# Mitigation of chytrid infection on tadpoles with the antimicrobial metabolites of *Xenorhabdus szentirmaii*

**DOI:** 10.1101/2024.08.27.609889

**Authors:** János Ujszegi, Zsófia Boros, Krisztián Harmos, Gábor Magos, Ábris Tóth, Judit Vörös, Andrea Kásler

**Affiliations:** Department of Evolutionary Ecology, HUN-REN Centre for Agricultural Research, Plant Protection Institute, H-1029, Budapest, Hungary; Department of Systematic Zoology and Ecology, Eötvös Loránd University, H-1117, Budapest, Hungary; Department of Genetics, Eötvös Loránd University, H-1117, Budapest, Hungary; Bükk National Park Directorate, H-3304, Eger, Hungary; Department of Zoology, University of Veterinary Medicine Budapest, H-1078, Budapest, Hungary; Amphibian and Reptile Conservation Group, MME Birdlife Hungary, H-1121, Budapest, Hungary; Doctoral School of Biology, Institute of Biology, Eötvös Loránd University, H-1117 Budapest, Hungary

**Keywords:** amphibian disinfection, EPN-EPB symbionts, frog-killing fungus, mitigation method, yellow-bellied toad

## Abstract

Survival of amphibian assemblages is threatened by many factors. Among them, chytridiomycosis, the disease caused by the chytrid fungus *Batrachochytrium dendrobatidis* (Bd) has great importance, also threatening the populations of the yellow-bellied toad (*Bombina variegata*), which has a scattered, isolated distribution in Hungary. Treatment with secondary metabolites of the entomopathogenic bacterium, *Xenorhabdus szentirmaii* is a promising method in the mitigation of chytridiomycosis, as recently demonstrated. To assess whether it may also be effective for lowering infection intensities in *B. variegata* tadpoles, we extracted cell-free culture media (CFCM) from liquid cultures of *X. szentirmaii* and tested their antifungal efficacy at a dilution of 0.1 % (v/v), while also measuring possible short-term malign effects on tadpoles experimentally infected with Bd or sham infection. According to our results, CFCM treatment alone did not compromise tadpoles’ survival probability, nor reduced the body mass and developmental speed of the individuals. At the same time, the treatment reduced the intensity and prevalence of Bd infection, but this effect was affected by the population of origin. The antimicrobial metabolites produced by *X. szentirmaii* may therefore be suitable for the safe mitigation of chytridiomycosis in *B. variegata* tadpoles, but further studies are needed, aiming to monitor and increase the efficacy of this method.

## 1. Introduction

Native amphibians are key taxa in many ecosystems serving as prey for predators (Burton & Likens 1975, Davic & Welsh 2004, Regester et al. 2006), consuming a wide array of invertebrates including pests and vector organisms (Khatiwada et al. 2016, Shuman-Goodier et al. 2019) and playing a weighty role in ecosystem services (Hocking & Babbitt 2014). Despite their significance, amphibians are going through a biodiversity crisis that started in the last century, becoming to be the most threatened vertebrate group in the present days (Monastersky 2014). The main causes of this decline are climate change, pollution, habitat loss, and emerging infectious diseases (Luedtke et al. 2023), which factors often exert a combined, synergistic negative effect on the populations of amphibian species (Koprivnikar 2010, Campbell Grant et al. 2016).

The natural defense mechanisms of amphibians are normally effective against a wide range of pathogens and parasites (Grogan et al., 2018), but pathogens introduced by human activities can have devastating effects on the naïve amphibian populations. Chytridiomycosis is the most serious infectious disease affecting amphibians (Scheele et al. 2019). The disease is caused by the chytrid fungi *Batrachochytrium dendrobatidis* (Bd) and *Batrachochytrium salamandrivorans* (Bsal), infecting the keratinous epidermal layers of the amphibian skin (Berger et al. 1998). These agents have already contributed to the decline or extinction of several hundred amphibian species and caused mass mortality events on all continents inhabited by amphibians (Van Rooij et al. 2015). Because Bsal has a much narrower distribution range (Lastra González et al. 2019), here we concentrate on the better-investigated and globally distributed Bd. The symptoms of heavy Bd infection are intensive sloughing or skin shedding, reddening on legs, and even ulcerations or skin lesions on ventral surfaces, ultimately loss of rightening reflex and body posture. The structural damage to the skin can impair skin breathing and osmoregulation, provoking shifts in electrolyte balance and finally leading to cardiac arrest (Voyles et al. 2009). In general, anuran tadpoles are less susceptible to the disease than in later life stages (Van Rooij et al. 2015), because keratinous elements are exhibited only in their mouthparts (Marantelli et al. 2004), thus they can act as reservoirs at natural habitats (Kilpatrick et al. 2010, Walker et al. 2010). However, the presence of Bd sometimes contributes to the loss of keratinized mouthpart structures, therefore leading to reduced feeding abilities and even lower survival (Blaustein et al. 2005). Several countermeasures against the disease have been proposed so far (Johnson et al. 2003, Woodhams et al. 2003, Harris et al. 2006, Woodward et al. 2014), but a widely applicable mitigation method against chytridiomycosis has not been found yet (Woodhams et al. 2012, Garner et al. 2016).

The addition or supplementation of mutualistic skin bacteria (bioaugmentation) associated to amphibians’ skin, that can prevent infections or disease propagation (Belden & Harris 2007, Krynak et al. 2016) is proposed to be a promising mitigation method against chytridiomycosis (Harris et al. 2009, Bletz et al. 2013, Rebollar et al. 2020), however, *in vivo* experiments usually reported moderate, or no mitigation effect against the disease (Woodhams et al. 2012, Küng et al. 2014, Rebollar et al. 2016). The target bacteria intended to settle on the new amphibian host can trigger an immune response (Rebollar et al. 2020), and may not establish or not produce the expected antifungal metabolites due to microbial competition or changing environmental factors experienced on the new host (Daskin et al. 2014, Robak & Richards-Zawacki 2018, Woodhams et al. 2018). Utilizing bacterial metabolites directly against Bd instead of trying to establish live cultures on amphibian hosts has also been tested (Bell et al. 2013, Madison et al. 2017, Ujszegi et al. 2023). This approach allows for avoiding most of the abovementioned problems, can be more safely controlled and the scope of the search for antifungal metabolites with broad-spectrum inhibition capabilities can be widened to cover microbial sources of non-amphibian origin too (Ujszegi et al. 2023).

Entomopathogenic nematode (EPN) species belonging to the *Steinernema* and *Heterorhabditis* genera are parasites of soil-dwelling insects (Akhurst 1982). Upon infection, EPN larvae release entomopathogenic bacteria (EPB) symbionts into the hemocoel of the insect host. These bacteria start to propagate and synthesize various secondary metabolites which suppress the insect’s immune response and accelerate its death. Furthermore, EPBs produce antimicrobial metabolites that protect the cadaver against microbial food competitors (Forst et al. 1997, Ogier et al. 2020). Since EPBs are culturable apart from their EPN hosts, these antimicrobial agents can be further utilized against plant-, livestock-, and human pathogens (Furgani et al. 2008, Böszörményi et al. 2009, Vozik et al. 2015, Wenski et al. 2020, Fodor et al. 2022). Moreover, secondary metabolites in cell-free culture medium (CFCM) extracted from liquid cultures of *Xenorhabdus szentirmaii* and *X. budapestensis* EPB species (Lengyel et al. 2005) have highly effective growth inhibition capabilities against Bd *in vitro*. In case of *X. szentirmaii*, CFCM can also be safely applied for significant reduction of Bd infection intensity on juvenile *Bufo bufo* individuals *in vivo* (Ujszegi et al. 2023). These results highlight the possibility of utilizing EPB metabolites in the mitigation of chytridiomycosis.

In Hungary, at least one Bd strain belonging to the Global Pandemic Lineage (GPL) is present (O’Hanlon et al. 2018) with the highest prevalence in frogs belonging to the *Bombina* and *Pelophylax* genera (Vörös et al. 2018). No mass mortalities have been observed so far, but yearly amphibian population surveys are focusing only on the breeding season. Therefore, mass die-offs during metamorphosis, when amphibians are the most susceptible to chytridiomycosis may remain hidden since their body size is small, and can decompose quickly. It could happen in the area of the Bükk National Park (Northern Hungary), where a recent study documented several incidents of lethal chytridiomycosis in the case of metamorphosing and juvenile *Bombina variegata* individuals (Harmos et al. 2021). Thus, to successfully combat Bd, the infection load should have been reduced already at the tadpole stage before individuals become more susceptible to the disease upon metamorphosis. In cooperation with the colleagues of the National Park, as a practical first step, we assessed whether *X. szentirmaii* CFCM may be effectively and safely applied for mitigation of infection intensity on *B. variegata* tadpoles experimentally exposed to Bd.

## 2. Materials & Methods

### 2.1. Culturing of bacteria

We prepared Luria broth agar (LBA) plates flooded with Luria broth (LB) (10 g casein peptone, 5 g yeast extract, 10 g sodium chloride, and 17 g agar dissolved in 1000 ml distilled water) as described previously (Ausubel et al. 1999). Indicator plates (LBTA) were supplemented with bromothymol blue and 2,3,5-Triphenyltetrazolium chloride and were used to distinguish antimicrobial peptide-producing (phase I) and non-producing (phase II) variants of *X. szentirmaii* (Leclerc & Boemare 1991). Fresh single-phase I colonies derived from frozen bacterial stocks were used for the experiment as previously described (Furgani et al. 2008, Böszörményi et al. 2009, Vozik et al. 2015). Microbiological media were obtained from Biolab Zrt. (Budapest, Hungary).

We cultured *X. szentirmaii* in liquid TGhLY medium (mTGhLY; 8 g tryptone, 2 g gelatine-hydrolysate, 4 g lactose, and 5 g yeast extract in 1000 ml distilled water) with a 7-day-old single colony grown on LBA. We adapted this method from an earlier study (Ujszegi et al. 2023) to provide optimal growth conditions in the same medium for both the Bd and the EPB with the addition of yeast extract. In all other respects, this medium is equivalent to the TGhL medium used for the culturing of Bd (see below). The EPB culture in this study was started with 5-10 ml of LB inoculated with a single colony of the respective bacterium picked from an LBTA indicator plate and incubated overnight at 28 °C in a water bath shaker (Lab-Line Orbital Shaker Water Bath, Marshall Scientific, USA). Each late-log phase inoculum was then added to 200 ml mTGhLY into 400 ml tissue culture flasks to create scale-up cultures.

### 2.2. Preparation of cell-free culture medium (CFCM)

We incubated scale-up cultures of *Xenorhabdus szentirmaii* for 7 days at 25 °C on an orbital shaker platform (Gallencamp, UK) With these preparation conditions, the production of antibiotic metabolites in *Xenorhabdus* cultures reaches a stationary phase in 5-6 days, containing the same amount of metabolites (Furgani et al. 2008, Böszörményi et al. 2009, Vozik et al. 2015). Then we centrifuged cultures at 6000 rpm for 20 minutes at 4 °C in 400 ml tubes using a JLA-10.500 type rotor (Avanti centrifuge J-26 XPI, Beckman Coulter, Indianapolis, USA). The supernatant was filtered through a sterile 0.22 µm nylon filter and centrifuged again at the same speed. We considered the resulting supernatant to be a cell-free culture medium (CFCM) of the antibiotic-producing *X. szentirmaii*. To confirm that CFCM was indeed cell-free, we diluted at least two replicates with sterile 2× LB, incubated them along with the experimental samples, and checked for bacterial growth on LBA plates. We stored CFCM at 4 °C in glass bottles until further use.

### 2.3. Experimental procedures

In May 2023 we collected 170 *Bombina variegata* eggs from puddles and wheel tracks at four localities (four distinct populations) in the Mátra mountains, Hungary (Haluskási út: 47.8866° N, 19.9938° E; Marháti út: 47.8911° N, 20.0418° E; Somhegy: 47.8868° N, 20.0076° E; Hidasi erdészház: 47.8831° N, 19.9834° E). We transported eggs to the Experimental Station Juliannamajor of the Plant Protection Institute, Centre for Agricultural Research located on the outskirts of Budapest (47.5479° N, 18.9349° E). We placed eggs of each of the four populations separately into plastic containers (32 × 22 × 16 cm) holding 0.7 l of reconstituted soft water (RSW; USEPA, 2002) at a constant temperature of 19.8 ± 0.4 °C and a light : dark cycle adjusted weekly to the conditions outside. Nine days after hatching, when all larvae reached development stage 25 (Gosner 1960), we started the experiment with 150 healthy-looking tadpoles (Haluskási út: N = 9 per treatment, 45 in total; Marháti út: N = 8 per treatment, 40 in total; Somhegy: N = 7 per treatment, 35 in total ; Hidasi erdészház: N = 6 per treatment, 30 in total). We reared tadpoles individually in opaque plastic boxes (17 × 12 × 9 cm) filled with 1 l RSW and fed them *ad libitum* with boiled and smashed spinach. Temperature in the laboratory was 18.2 ± 0.3 °C (mean ± SD) during the experiment. The light : dark cycle was adjusted weekly to outdoor conditions. We assigned tadpoles to five treatments (Table 1), using stratified randomization considering their population of origin: we exposed them to RSW (Treat 1), sterile TGhLY medium (Treat 2 and 3), or liquid culture of Bd (Treat 4 and 5; please see below for details) in the next 22 days. All hatchlings beyond those 150 that were arranged into the five treatments, were infected with Bd and kept the same way to increase the sample size of reference for initial Bd infection (see below). We arranged rearing boxes randomly on the laboratory shelves and changed water twice a week using different dip nets for each treatment to prevent cross-contamination. We monitored survival daily and noted any incidence of death.

**Table 1.** Treatment combinations.

|  | medium | Bd exposure | CFCM treatment |
| --- | --- | --- | --- |
| Treat 1 | no | no | no |
| Treat 2 | yes | no | no |
| Treat 3 | yes | no | yes |
| Treat 4 | yes | yes | no |
| Treat 5 | yes | yes | yes |

Twenty-two days after the start of the experiment, we randomly selected five individuals from each treatment group, and humanely euthanized them together with all surplus individuals in a water bath containing 6.6 g l^-1^ tricaine-methanesulfonate (MS-222) buffered to neutral pH with the same amount of Na_2_HPO_4_. We conserved dead individuals in 96 % EtOH and stored samples at 4 °C until further analysis as a reference for the initial Bd infection status of the animals before mitigation. We changed the boxes of the remaining individuals and started mitigation treatments by exposing them to either RSW, mTGhLY, or CFCM at a dilution of 0.1 % (v/v) (Table1). We constantly treated animals for another 22 days re-administering treatments after each water change. On the last day of treatments, we gently blotted and weighed individuals to the nearest mg (OHAUS-PA213 analytical balance, Ohaus Europe Gmb, Nanikon, Switzerland), euthanized and preserved them as described above.

### 2.4. Maintaining Bd cultures and experimental exposure

We used the global pandemic lineage (GPL) of Bd. The isolate (Hung_2014) originated from a *B. variegata* collected alive in 2014 by J. Vörös in the Bakony Mountains, Hungary, and isolated by M.C. Fisher and colleagues (Imperial College London, London, UK). We maintained parallel cultures in TGhL medium (mTGhL; 8 g tryptone, 2 g gelatine-hydrolysate, and 4 g lactose in 1000 ml distilled water) in 25 cm^2^ cell culture flasks at 4 °C and passaged them every three months into sterile mTGhL.

One week before performing experimental infections, we inoculated 100 ml mTGhLY with 2 ml of Bd stock culture in a 175 cm^2^ cell culture flask and incubated it for seven days at 21 °C. We assessed the concentration of intact zoospores using a Bürker chamber at ×400 magnification before every inoculation. During inoculation of tadpoles’ rearing boxes, the mean initial concentrations were ∼ 7.5 × 10^5^ zoospores (zsp) ml^-1^ in the flasks. After each water change, we inoculated 1 ml of these cultures into the tadpoles’ rearing boxes holding 1 l RSW, resulting in ∼ 750 zsp ml^-1^ concentration in the rearing boxes. We inoculated controls with the same quantity of sterile mTGhLY or RSW according to the treatments. Contaminated water and equipment were disinfected overnight with VirkonS before disposal (Johnson et al. 2003).

### 2.5. Assessment of infection intensity

We cut out, then homogenized the whole mouthparts of the preserved tadpoles, extracted DNA from samples using PrepMan Ultra Sample Preparation Reagent (Thermo Fisher Scientific, Waltham, Massachusetts, USA) according to previous recommendations (Boyle et al. 2004), and stored extracted DNA at -20 °C until further analyses. We assessed infection intensity using real-time quantitative polymerase chain reaction (qPCR) following a standard amplification methodology targeting the ITS-1/5.8S rDNA region (Boyle et al. 2004) on a BioRad CFX96 Touch Real-Time PCR System (BioRad Laboratories, Hercules, USA). To avoid PCR inhibition by ingredients of PrepMan, we diluted samples ten-fold with double-distilled water. We ran samples in duplicate, and in case of equivocal results, we repeated reactions in duplicate. If this again returned an equivocal result, we considered the sample to be Bd positive (Kriger et al. 2006). Genomic equivalent (GE) values were estimated from standard curves based on five dilutions of a standard (1000, 100, 10, 1, and 0.1 zoospore GE; provided by J. Bosch; CSIC-University of Oviedo, Spain).

### 2.6. Statistical analyses

From the analyses, we excluded all the reference individuals preserved for inspecting initial Bd infection status, and a further individual from the infection analyses which started metamorphosis before the end of the experiment. We assessed treatment effects on survival, the stage of larval development, body mass, Bd prevalence, and infection intensity. For each dependent variable, we ran a model (see model specifications below) with treatment and population of origin (population hereafter) as categorical fixed factors and their interaction. For the analysis of survival, we used Cox’s proportional hazards model (R package ‘coxme’) with treatment as the explanatory variable. The structure of the data did not allow for including population as another fixed factor. We entered the number of days until death as the dependent variable and individuals that survived until the termination of the experiment were treated as censored observations. To analyse variation in the stage of larval development and body mass, we used general linear models (LM; ‘lm’ function of the ‘nlme’ package). In the analysis of body mass, we further included developmental stage as a covariate. To analyse Bd prevalence, we used generalized linear models (‘glm’ function of the ‘nlme’ package) with binomial distribution and logit link function. For the investigation of infection intensity, we averaged GE values obtained from qPCR runs for each sample and analysed resulting estimates corrected for body mass using generalized linear mixed models (GLMM) with negative binomial distribution and a log link function using the ‘glmmTMB’ package (Brooks et al. 2017). All tests were two-tailed, and we checked model fits in the case of all dependent variables by visual inspection of diagnostic plots. We applied a backward stepwise model simplification procedure (Grafen & Hails 2002) to avoid potential problems due to the inclusion of non-significant terms (Engqvist 2005). We used the significance value of alpha = 0.05. In case of significant effect by any grouping explanatory variable, we applied post-hoc tests by calculating pre-planned linear contrasts (Ruxton & Beauchamp 2008), correcting the significance threshold for multiple testing using the false discovery rate (FDR) method (Pike 2011). All analyses were conducted in ‘R’ (version 4.0.5).

## 3. Results

Survival was not affected by any of the treatments (Cox model: z = -1.24, p = 0.21; Fig 1). The stage of larval development was significantly affected by treatment (LM: F_4,94_ = 17.19, p < 0.001; Fig 2A) and by population (F_3,94_ = 4.27, p < 0.001), but not their interaction (F_12,82_ = 0.58, p = 0.86). Pairwise comparisons revealed that developmental speed was significantly lower in the group Treat 1 compared with all other groups, but Bd exposure and / or CFCM treatment did not affect it (Table 2). Body mass of the tadpoles at the end of the experiment was not affected by population (LM: F_3,92_ = 1.41, p = 0.25) or its interaction with treatment (LM: F_12,80_ = 1.01, p = 0.45), but it was significantly affected by treatment alone (F_4,95_ = 35.15, p < 0.001; Fig 2B) and developmental stage (LM: F_1,95_ = 42.87, p < 0.001). Pairwise comparisons revealed that Treat 1 significantly reduced body mass compared to all other treatment groups. However, Bd exposure or treatment with CFCM did not affect body mass (Table 3, Fig 2B).

**Figure 1:**
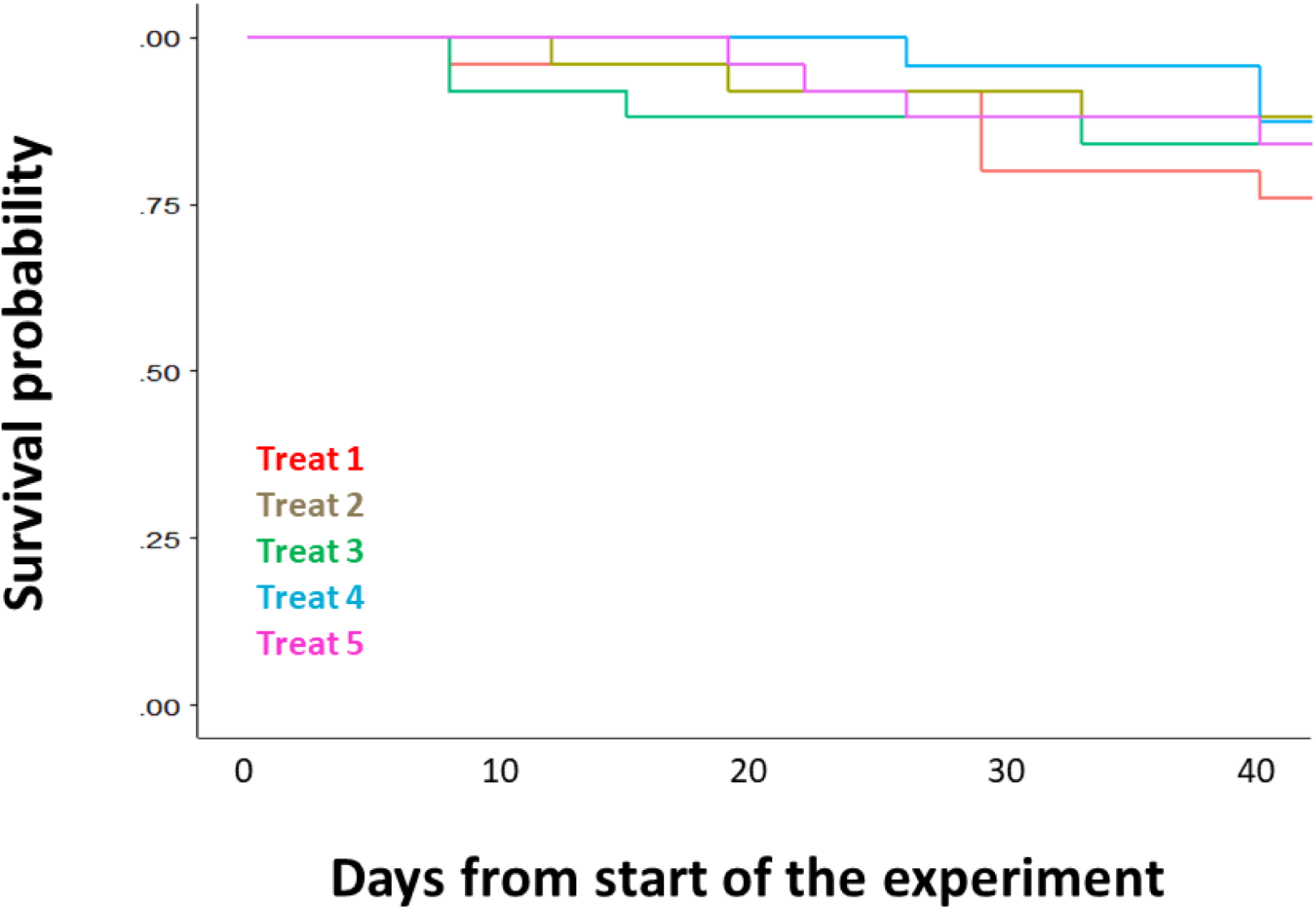
Survival of *Bombina variegata* tadpoles over the experiment in the five treatments. For explanation of treatments please see Table 1.

**Figure 2:**
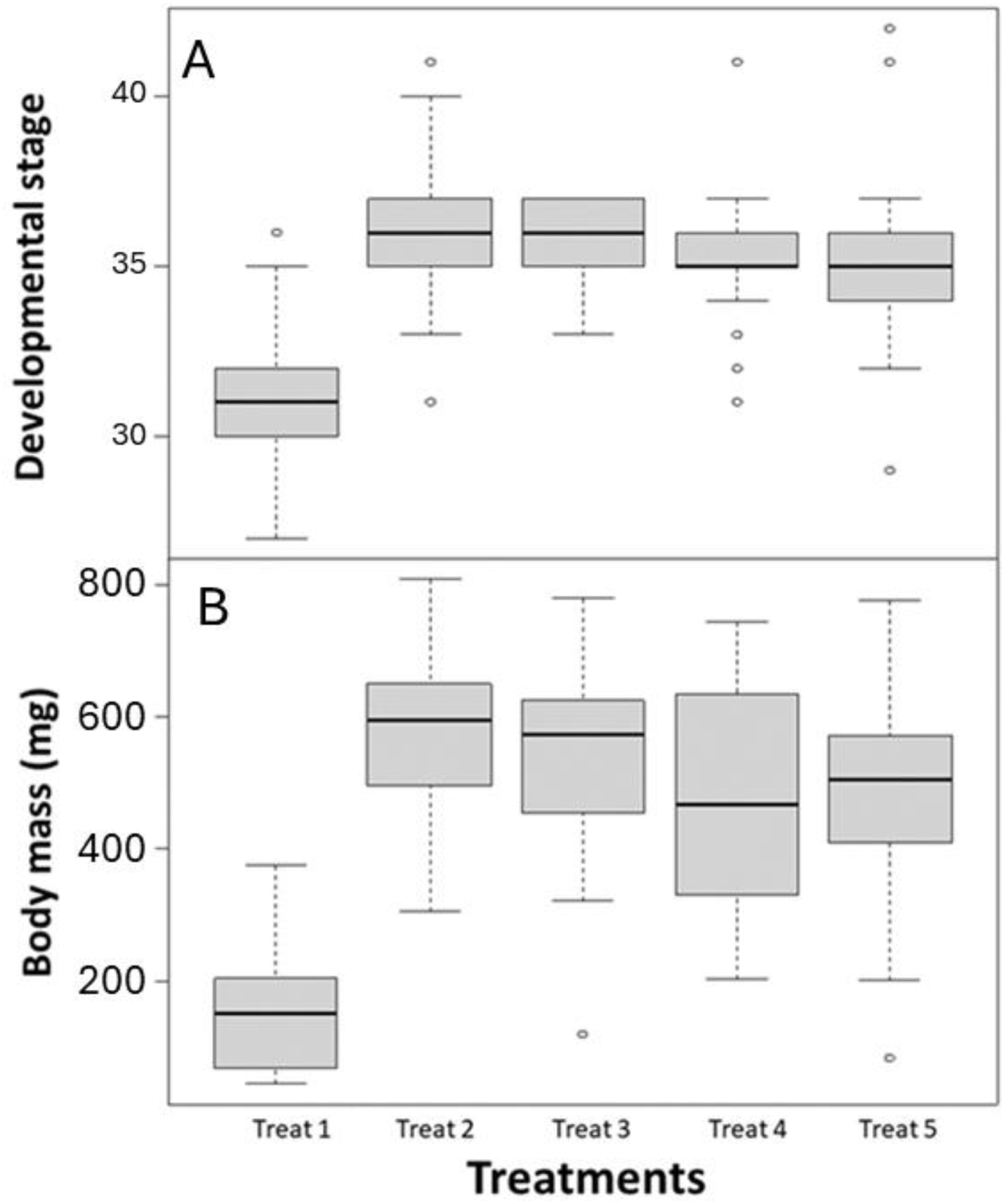
Responses of *Bombina variegata* tadpoles to the five treatments in terms of the (A) developmental stage (Gosner) and (B) body mass at the termination of the experiment. In boxplots, horizontal lines and boxes represent medians and interquartile ranges (IQR), respectively, while whiskers extend to IQR ± 1.5× IQR and dots indicate more extreme data points. For explanation of treatments please see Table 1.

**Table 2.**
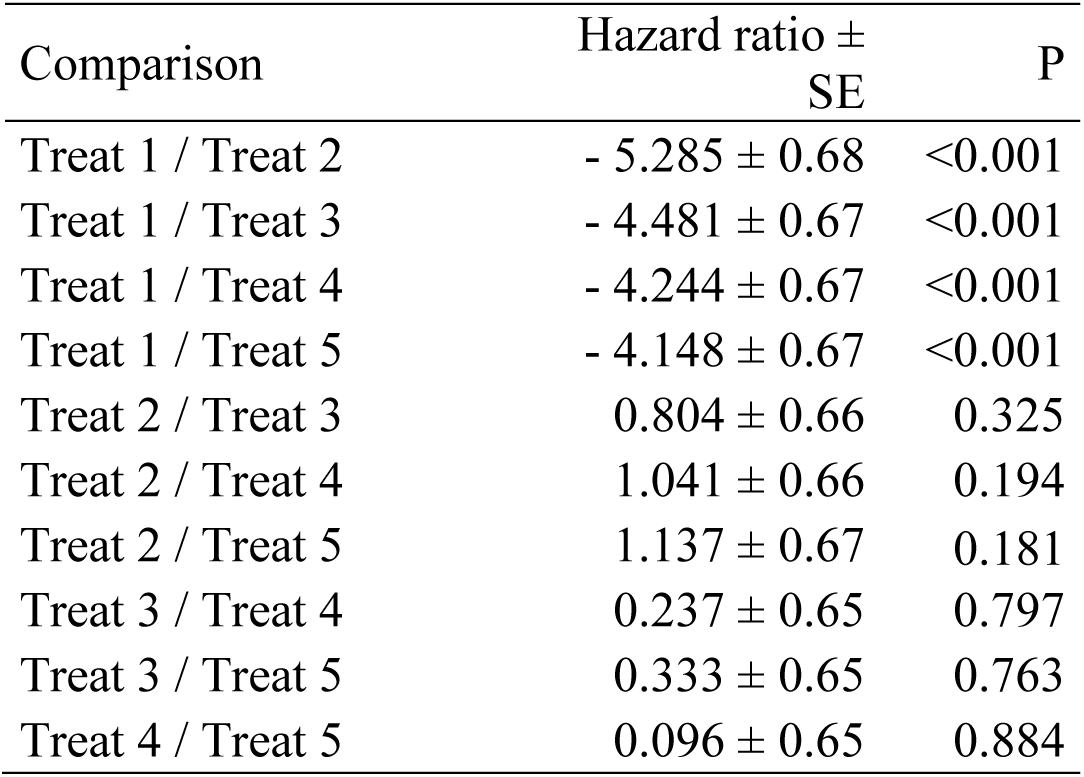
Pairwise comparisons for the effects of treatments on developmental stage.

| Comparison | Hazard ratio ±<br>SE | P |
| --- | --- | --- |
| Treat 1 / Treat 2 | - 5.285 ± 0.68 | <0.001 |
| Treat 1 / Treat 3 | - 4.481 ± 0.67 | <0.001 |
| Treat 1 / Treat 4 | - 4.244 ± 0.67 | <0.001 |
| Treat 1 / Treat 5 | - 4.148 ± 0.67 | <0.001 |
| Treat 2 / Treat 3 | 0.804 ± 0.66 | 0.325 |
| Treat 2 / Treat 4 | 1.041 ± 0.66 | 0.194 |
| Treat 2 / Treat 5 | 1.137 ± 0.67 | 0.181 |
| Treat 3 / Treat 4 | 0.237 ± 0.65 | 0.797 |
| Treat 3 / Treat 5 | 0.333 ± 0.65 | 0.763 |
| Treat 4 / Treat 5 | 0.096 ± 0.65 | 0.884 |

**Table 3.** Pairwise comparisons for the effects of treatments on body mass.

| Comparison | Hazard ratio ±<br>SE | P |
| --- | --- | --- |
| Treat 1 / Treat 2 | - 222.35 ± 49.8 | <0.001 |
| Treat 1 / Treat 3 | - 224.35 ± 46.8 | <0.001 |
| Treat 1 / Treat 4 | - 169.24 ± 46.3 | 0.001 |
| Treat 1 / Treat 5 | - 167.00 ± 46.3 | 0.001 |
| Treat 2 / Treat 3 | - 2.25 ± 38.5 | 0.954 |
| Treat 2 / Treat 4 | 53.11 ± 38.6 | 0.216 |
| Treat 2 / Treat 5 | 55.35 ± 38.6 | 0.216 |
| Treat 3 / Treat 4 | 55.36 ± 37.8 | 0.216 |
| Treat 3 / Treat 5 | 57.60 ± 37.8 | 0.216 |
| Treat 4 / Treat 5 | 2.24 ± 37.8 | 0.954 |

Since all tested individuals from uninfected treatment groups remained Bd negative, we can exclude the possibility of cross-contamination. Bd prevalence was 71 % in experimentally infected individuals chosen as reference for initial Bd infection right before the start of the CFCM treatment, with an average infection intensity of 14.7 (0.0-111.8) GE (median and interquartile range) for all individuals. CFCM treatment resulted in significantly lower Bd prevalence (GLM: z = -2.83, p = 0.009; 54 % vs. 89 % in the non-treated group) and infection intensity (z = -3.65, p < 0.001; 0.14 [0.0-1.18] GE/gram body weight [GE/gbw; median and interquartile range]; Fig 3) compared to the Bd exposed group without CFCM treatment (0.59 [0.13-1.87] GE gbw^-1^ [median and interquartile range]; Fig 3). However, the population also affected this antifungal effect since its interaction with treatment was also significant in the case of infection intensity (z = 3.58, p < 0.001) and marginally non-significant in the case of Bd prevalence (z = 1.83, p = 0.067). Omitting this interaction, the effect of population alone on Bd prevalence was significant (z = 2.29, p = 0.022).

**Figure 3:**
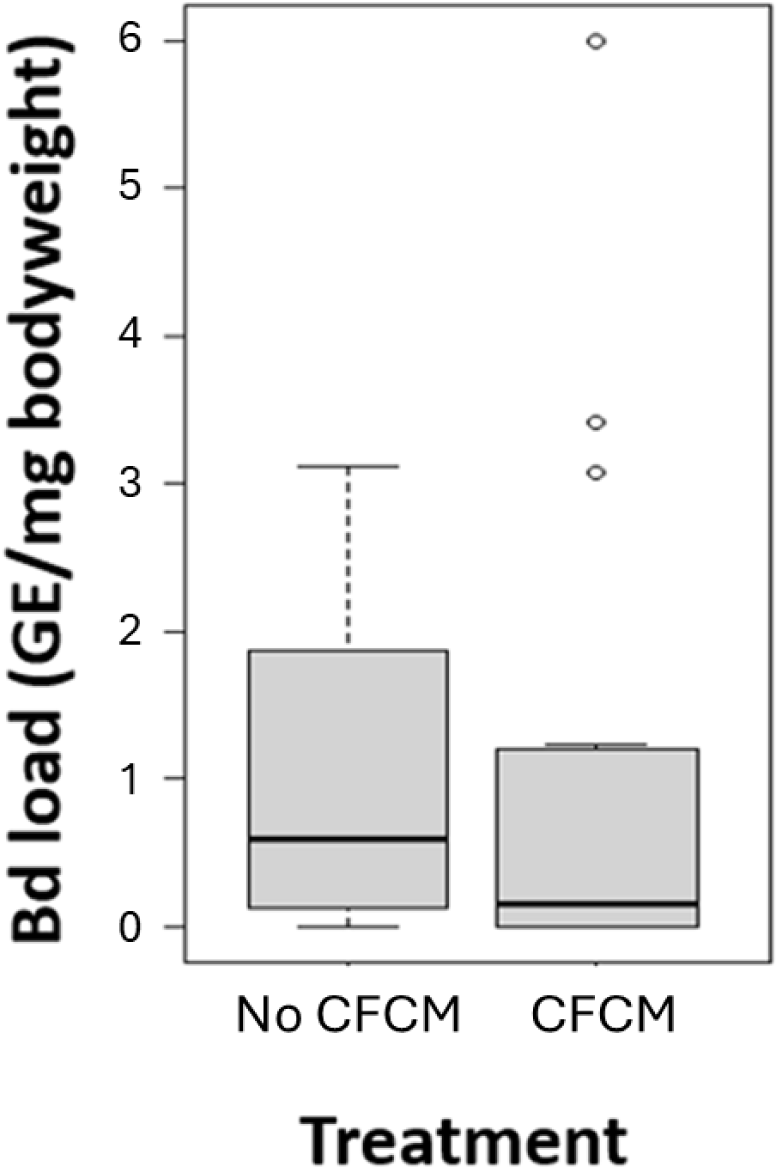
Tadpoles’ infection intensities from the two Bd exposed groups (Treat 4 and Treat5) in the absence and the presence of 0.1 % (v/v) cell-free culture medium (CFCM) treatment. Horizontal lines and boxes represent medians and interquartile ranges (IQR), respectively, while whiskers extend to IQR ± 1.5× IQR and dots indicate more extreme data points.

## 4. Discussion

Treatment with CFCM of the *X. szentirmaii* EPB had no short term negative effect on the measured life history traits of *Bombina variegata* tadpoles, similarly to former results on juvenile *Bufo bufo* individuals exposed to much more concentrated CFCM solutions (Ujszegi et al. 2023). Moreover, microbial medium at the applied dilution seemed to exert a beneficial effect in the terms of developmental speed and body mass of the tadpoles, compared with conspecifics reared in pure RSW throughout the experiment. This is likely due to the absorption of extra nutritional compounds from the media through their gut, which accelerated tadpoles’ growth and development. However, whether CFCM treatment has other negative effects on the long run for the treated amphibians, for example through the alteration of skin microbiome, is still need to be investigated in future studies.

Prevalence of chytridiomycosis can reach high values in *B. variegata* at several natural habitats across its whole distribution (Sztatecsny & Glaser 2011, Baláž et al. 2014, Kolenda et al. 2017, Vojar et al. 2017, Vörös et al. 2018), likely contributing to local population declines (Canestrelli et al. 2013). Additionally, mortality events that have been documented (Baláž et al. 2014, Harmos et al. 2021) further emphasize that intervention may be necessary soon to prevent populations from catastrophic losses. We demonstrated, that *X. szentirmaii* CFCM solution applied at a concentration as low as 0.1 % v/v is still able to inhibit Bd growth *in vivo* in *B. variegata* tadpoles. This extreme effectiveness is in line with former findings, where dilution close to 0.1 % CFCM of the same microbe still showed *in vitro* growth inhibition of Bd (Ujszegi et al. 2023). However, growth inhibition capabilities varied between the host’s populations of origin. Specimens of *B. variegata* secrete potent AMPs in their skin (Mignogna et al. 1993), and intraspecific differences in AMP synthesis can reflect the differences of populations in the sensitivity to chytridiomycosis in other amphibians (Tennessen et al. 2009, Bradley et al. 2015). Therefore different *B. variegata* populations may differ in their immunity and skin-secreted AMP repertoire, which affect their tolerance of Bd infection, and probably differently interact with antimicrobial metabolites in the CFCM. Also, distinct populations may harbor different skin microbiome compositions (Woodhams et al. 2014) variously affecting, and interacting with the beneficial effect of the CFCM treatment. Improvement of the method’s efficacy to reliably achieve a more uniform effect could help to remove this population effect very likely.

One possible option for enhancing effectivity would be the application of a more concentrated CFCM solution, but this can be problematic upon constantly treating tadpoles, exposing them to CFCM through their ambient water. Since CFCM is a culture medium, its ingredients can promote excessive bacterial bloom in the water already at the dilution of 0.5 % v/v, which compromises dissolved oxygen level, threatening the tadpoles’ health (Ujszegi et al. unpublished). Furthermore, CFCM concentrated more than 15% can be harmful to individuals directly (Ujszegi et al. 2023) thus taken together, this direction to enhance the effectiveness of CFCM treatment on amphibians is not recommended. An alternative option could be the treatment of tadpoles in separate enclosures out of their rearing water for a limited time on several consecutive occasions to reduce pathogen load or clear infection. In this case, since tadpoles are exposed to CFCM in fresh water on every occasion and only for a limited time, more concentrated CFCM could be used to enhance effectivity, such as in the case of juvenile *B. bufo* individuals (Ujszegi et al. 2023). This direction deserves more detailed research and could be a very useful method in the laboratory, or in captive breeding facilities where disinfection is crucial for preventing disease outbreaks (Kueneman et al. 2022), even in the case of *B. variegata* (Stagni et al. 2004). However, for such CFCM treatments, all necessary instruments are better available indoors, and the number of individuals is more manageable than in the field, therefore *in situ* application of this direction is very limited.

Since a low level of infection usually causes no clinical signs or mortality (Vredenburg et al. 2010, Cheng et al. 2011) and amphibian populations can coexist with Bd (Rowley & Alford 2013), a complete clearance of infection may not be essential. It is possible, that the efficiency achieved by the treatment with 0.1 % *X. szentirmaii* CFCM can even be sufficient for the mitigation of chytridiomycosis at *B. variegata* habitats, because co-existence with enzootic Bd may lead to immunization (Ramsey et al. 2010, McMahon et al. 2014) and allow for adaptation to the disease via the spread of resistance alleles (Bataille et al. 2015, Savage & Zamudio 2016, Voyles et al. 2018) or alternative reproductive strategies (Spitzen-Van Der Sluijs et al. 2017) in the populations. Whether the observed effect is sufficient for halting mortality events by Bd in natural populations, and whether the addition of CFCM would be harmful to other aquatic organisms besides amphibians (such as pond vegetation and macroinvertebrates) at the habitats should be long-term monitored. Given that CFCM has got wide array of antimicrobial activity, and the media contains many extra nutrients, investigating whether the treatment causes long-term shifts in the aquatic microbial composition and therefore in the flow of energy and nutrients would be also important before routinely applying this method at natural habitats.

Since a universally applicable mitigation method against chytridiomycosis for most of the host species with the same level of efficacy won’t be found very likely (Garner et al. 2016), researchers and nature conservation specialists must focus their effort on species that are endangered, or largely affected by chytridiomycosis. Scientists should find locally adaptable and successful *in situ* mitigation methods designed for the focal species of conservation (e.g. Thumsová et al., 2024), considering their special needs and characteristics. Going back to *B. variegata*, these frogs mostly live in small water bodies and wheel track puddles. In such environments, the desired CFCM amount (0.1 % v/v) could be easily achieved needing no more than some litres for the treatment of the whole water body. CFCM can still be produced quickly and cheaply in such quantities. Furthermore, due to its extreme thermostability, CFCM does not require special handling during transportation and would last for a long time after application. Based on these characteristics and our results, improving this method to be applied for *in situ* mitigation purposes to preserve *B. variegata* populations would be worthwhile in the future.

## Acknowledgments

We are thankful to A. Hettyey, Z. Mikó, N. Ujhegyi, D. Herczeg, and M. Szederkényi for their assistance and help during the experiment, A. Hettyey for providing the institutional background for the experiment, V. Bókony for help in statistics and M. Z. Németh, and all colleagues from the Department of the Plant Pathology for allowing us to use their lab facilities. T. Vellai provided us with the laboratory capacity, and A. Fodor helped with professional guidance at the Department of Genetics, Eötvös Loránd University. Zoospore genomic equivalent standards were kindly offered by J. Bosch, and Bd isolate was sent by M. C. Fisher. The research was supported by the Lendület Programme of the Hungarian Academy of Sciences (MTA, LP2012-24/2012) and the New National Excellence Program of the Ministry for Innovation and Technology from the source of the National Research, Development, and Innovation Fund (ÚNKP-23-4 for U.J., ÚNKP-22-3 and ÚNKP-23-4 for A. K., ÚNKP-23-6 for Á. T.). Open access funding was provided by Eötvös Loránd University. The Ethical Commission of the Plant Protection Institute approved experimental procedures and research was carried out according to the permits issued by the Government Agency of Pest County (PEI/001/1797-3/2015, PE/EA/00270-6/2023) and the Government Agency of Heves County (HE/TVO/00022-10/2023). The authors have no conflict of interest to declare.

## Notes

### Competing Interest Statement

The authors have declared no competing interest.

